# Widespread FUS mislocalization is a molecular hallmark of ALS

**DOI:** 10.1101/491787

**Authors:** Giulia E. Tyzack, Raphaelle Luisier, Doaa M. Taha, Jacob Neeves, Miha Modic, Jamie S. Mitchell, Ione Meyer, Linda Greensmith, Jia Newcombe, Jernej Ule, Nicholas M. Luscombe, Rickie Patani

**Author notes:** These authors contributed equally to this work (Giulia E. Tyzack, Raphaelle Luisier). Correspondence should be addressed to Nicholas M. Luscombe and Rickie Patani.

## Abstract

Amyotrophic lateral sclerosis (ALS)-causing mutations clearly implicate ubiquitously expressed and predominantly nuclear RNA binding proteins (RBPs), which form pathological cytoplasmic inclusions in this context. However, the possibility that wild-type RBPs mislocalize without necessarily becoming constituents of ALS cytoplasmic inclusions themselves remains unexplored. We hypothesized that nuclear-to-cytoplasmic mislocalization of the RBP Fused in Sarcoma (FUS), in an unaggregated state, may occur more widely in ALS that previously recognized. To address this hypothesis, we analysed motor neurons (MNs) from an human ALS induced-pluripotent stem cells (iPSC) model caused by the VCP mutation. Additionally, we examined mouse transgenic models and post-mortem tissue from human sporadic ALS cases. We report nuclear-to-cytoplasmic mislocalization of FUS in both VCP-mutation related ALS and, crucially, in sporadic ALS spinal cord tissue from multiple cases. Furthermore, we provide evidence that FUS protein binds to an aberrantly retained intron within the SFPQ transcript, which is exported from the nucleus into the cytoplasm. Collectively, these data support a model for ALS pathogenesis whereby aberrant intron-retention in SFPQ transcripts contributes to FUS mislocalization through their direct interaction and nuclear export. In summary, we report widespread mislocalization of the FUS protein in ALS and propose a putative underlying mechanism for this process.

## Introduction

Amyotrophic lateral sclerosis (ALS) is a relentlessly progressive neurodegenerative condition, which remains incurable due to our incomplete understanding of the molecular pathogenesis. Genetic discoveries in ALS strongly implicate ubiquitously expressed regulators of RNA-processing (Taylor *et al.,* 2016). Pathologically, in 97% of cases TDP-43 protein is mislocalized from the nucleus to the cytoplasm, where it aggregates (Neumann *et al.,* 2006). However, ALS-causing mutations in Fused in Sarcoma (FUS) and superoxide dismutase (SOD1) conspicuously lack TDP-43 proteinopathy in the majority of cases (Mackenzie *et al.,* 2007). FUS aggregation is a recognized feature of FUS mutation-related ALS (Vance *et al.,* 2009). However, the concept that wild-type FUS nuclear-to-cytoplasmic mislocalisation - rather than aggregation - might be a more widespread feature of other forms of ALS has not been systematically assessed to our knowledge. Indeed such mislocalisation may have evaded detection thus far due to bias towards studying aggregates/inclusions rather than perturbed subcellular localisation of unaggregated proteins alone. This possibility is strengthened by the facts that i) impaired nuclear-cytoplasmic compartmentalization is increasingly recognized as a key feature of ALS (Boeynaems *et al.,* 2016) and ii) FUS is known to shuttle between the nucleus and cytoplasm.

Here we systematically investigated FUS protein localisation across a human iPSC model, mouse transgenic models and human post-mortem tissue from multiple cases of sporadic ALS. We find that the nuclear-to-cytoplasmic mislocalisation of FUS is a more widespread feature of ALS than previously recognized. Furthermore, we present evidence that supports a putative molecular mechanism for this mislocalisation through interaction between FUS and the aberrant intron retaining SFPQ transcript.

## Results

### Nuclear-to-cytoplasmic mislocalisation of FUS in human and mouse VCP-mutant ALS models

We have previously reported robust differentiation of human induced-pluripotent stem cells (hiPSC) into highly enriched (>85%) cultures of comprehensively validated and functionally characterized spinal cord motor neurons (Hall *et al.,* 2017; Maffioletti *et al.,* 2018). Using this approach, we have found clear molecular pathogenic phenotypes in a human iPSC model of VCP-related ALS (4 clones from 2 patients with the following mutations: VCP^155C^ and VCP^R191Q^) (Hall *et al.,* 2017; Luisier *et al.,* 2018). We harnessed this well characterized ALS model to investigate the subcellular localisation of FUS protein, which revealed a modest but statistically significant (P < 0.001) decrease in nuclear-to-cytoplasmic localisation during motor neuron differentiation (Figure 1A). We next sought to validate this finding *in vivo* by examining tissue sections from a mouse VCP transgenic model (VCP^A232E^). Additionally, we examined tissue from the SOD1^*G93A*^ mouse model as a comparator. Using linear mixed model analysis that accounts for inter-animal variation we showed that although FUS remained within the nucleus in the SOD1 mouse model (χ^2^ (1)=0.031, p=0.955), nuclear-to-cytoplasmic mislocalisation abounded in the VCP mouse model, with a reduction in FUS nuclear to cytoplasmic ratio of −4.0176 ± 1.1775 (χ^2^ (1)=8.164, p=0.0042) (Figure 1B).

**Figure 1.**
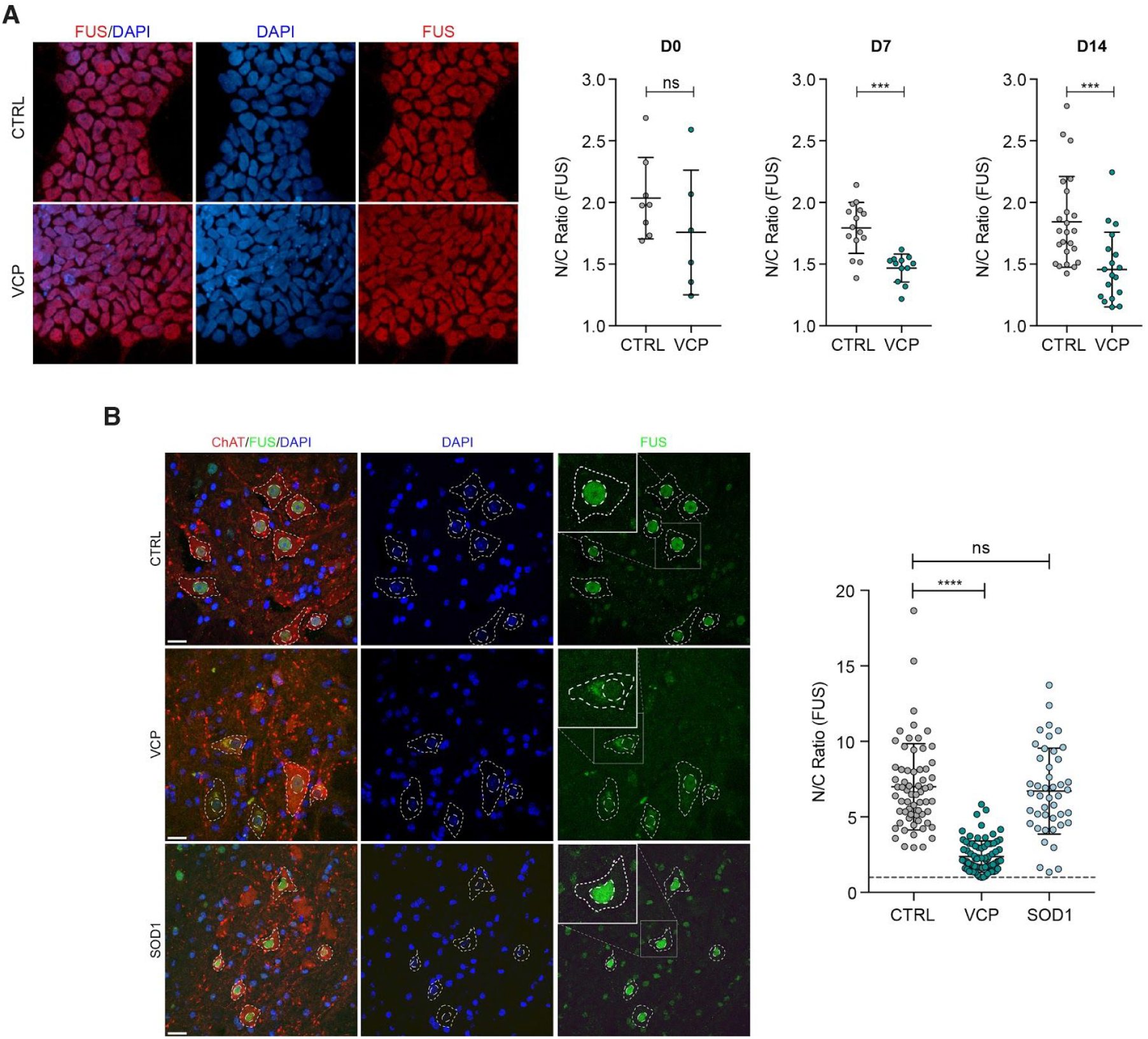
Nuclear-to-cytoplasmic mislocalisation of FUS in human and mouse VCP-mutant ALS models. **(A)** Subcellular localization of FUS determined by immunocytochemistry in iPSCs, neural precursors (NPCs) and patterned motor neuron precursors (pMNs). The ratio of the average intensity of the FUS immunolabelling in the nucleus (N) vs cytoplasm (C) was automatically determined in both CTRL and VCPmucells. Data shown is average N/C ratio (± s.d.) per field of view from 4 control and 4 VCPmulines.P value from unpaired t-test with Welch’s correction. **(B)** Analysis of the subcellular localisation of FUS in MNs in the ventral spinal cord of wild-type, *VCP^A232E^* and *SOD1^G93A^* mice. MN cytoplasm was identified by ChAT staining, nuclei were counterstained with DAPI. Images were acquired as confocal z-stacks using a Zeiss 710 confocal microscope with a z step of 1μm, processed to obtain a maximum intensity projection and analysed using Fiji. MNs were identified by ChAT staining, and the nuclear and cytoplasmic area were manually drawn based on DAPI and ChAT staining respectively. For each region of interest, the average FUS immunoreactivity intensity was measured, background was subtracted, and the ratio between nuclear and cytoplasmic average intensity was calculated. Data shown is nuclear/cytoplasmic (N/C) ratio (mean± s.d.) per cell from 4 wild-type, 4 *SOD1^G93A^* and 3 *VCP^A232E^* mice. Scale bar: 20μm. Linear mixed effects analysis of the relationship between FUS localisation and either VCP or SOD1 mutation to account for idiosyncratic variation due to animal differences: SOD1 mutation does not affect FUS localisation (χ^2^ (1)=0.6387, p=0.4242), lowering it by about −0.8152 ± 1.1152 (standard errors), whilst VCP mutation does affect FUS localisation (χ^2^ (1)=8.3145, p=0.003933), lowering it by −4.2175 ± 0.9731 (standard errors). Putative FUS inclusions are indicated in the VCP condition by white arrowheads.

### Nuclear-to-cytoplasmic mislocalisation of FUS in human sporadic ALS

Having established that FUS was mislocalized in VCP-related ALS models, but not in SOD1, we next sought to address the generalizability of this finding across sporadic forms of ALS (which represent 90% of all cases). To this end, we examined post-mortem spinal cord tissue from 12 sporadic ALS cases and 8 healthy controls (Figure 2A). We found clear evidence of nuclear-to-cytoplasmic FUS mislocalisation in these sporadic ALS cases, but *in the absence of cytoplasmic FUS inclusions*.

**Figure 2.**
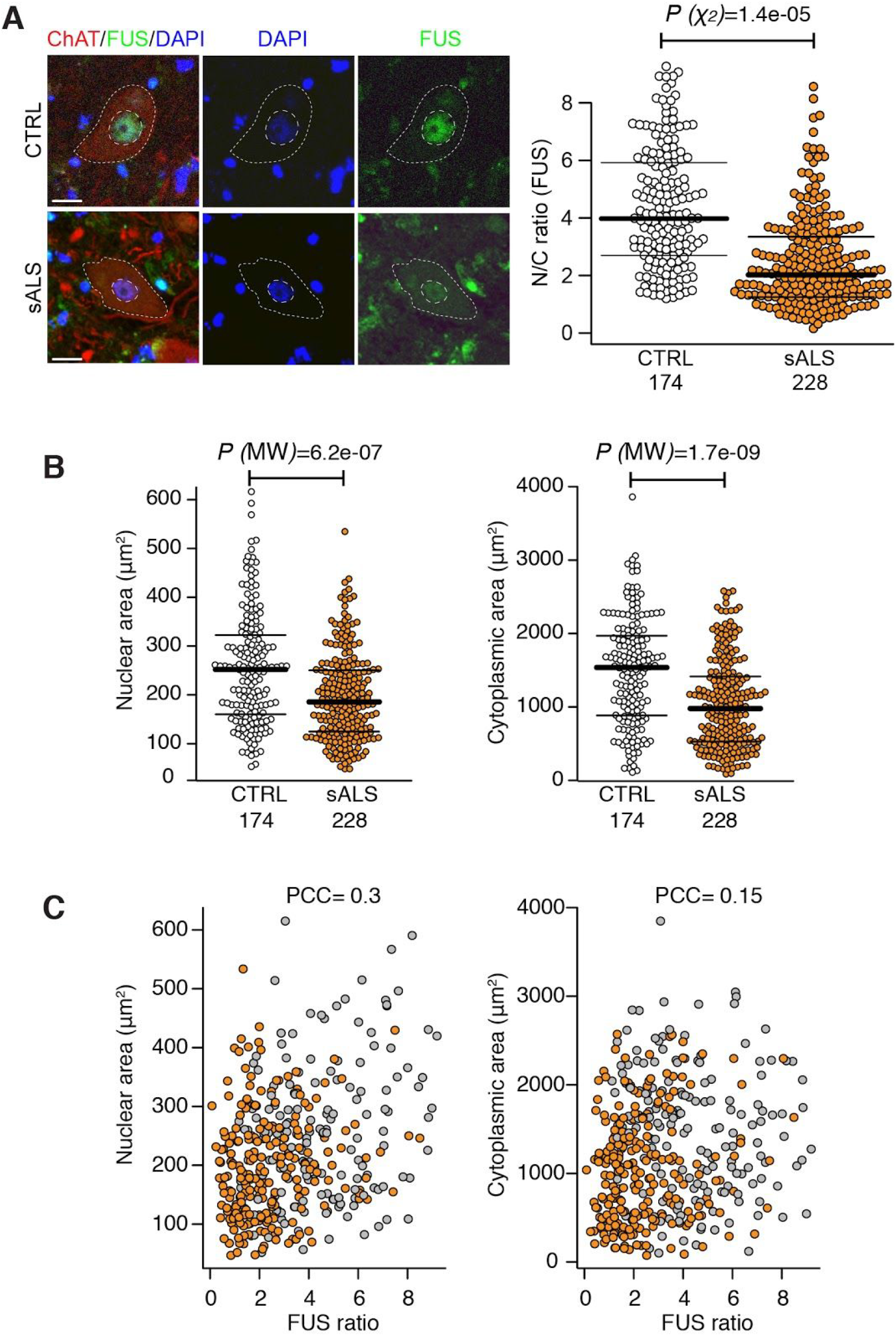
Nuclear-to-cytoplasmic mislocalisation of FUS in human sporadic ALS. **(A)** Analysis of the subcellular localisation of FUS in MNs in the ventral spinal cord of healthy controls (n=8) and patients with sporadic ALS (sALS) (n=12). MN cytoplasm was identified by ChAT staining, nuclei were counterstained with DAPI. N/C ratio of FUS immunoreactivity was measured as described in Figure 1B. Only MNs with a visible nucleus were considered for the analysis. We used R (R Core Team, 2012) and lme4 (Bates *et al.,* 2015) to perform a linear mixed effects analysis of the relationship between FUS localisation and sALS disease. Scale bar 20μm. Data shown is N/C ratio (mean ± s.d.) per cell from at least 8 cases per group. **(B)** Nuclear and cytoplasmic areas of MN nuclei and cytoplasm in sALS (n=12) vs control (n=8). Data shown is mean ± s.d. *P*-values from non-parametric Mann-Whitney’s test. **(C)** Single-cell measurement of nuclear (right) and cytoplasmic area (left) shows no correlation with FUS nuclear to cytoplasmic ratio as shown with scatter plots. PCC=Pearson correlation coefficient.

A reduction in motor neuronal soma size has been previously shown in an in vitro model of ALS which associates with increase apoptosis (Kiskinis *et al.,* 2014). We thus investigated whether changes in FUS subcellular localisation in sALS samples accompany alterations in cell morphology. Using single-cell analysis of nuclear and cell size, we demonstrated significant reductions in both nuclear and cytoplasmic areas in sALS vs control MNs (Figure 2B) which did not correlate with reduction in FUS nuclear-to-cytoplasmic localisation in sALS cases (Figure 2C).

### FUS binds to an aberrantly retained intron within the SFPQ transcript, which is exported from the nucleus

Having found clear evidence that FUS nuclear-to-cytoplasmic mislocalisation is more widespread in ALS than previously recognized, we sought to understand it’s molecular interplay with 167 aberrant intron retaining transcripts that we recently described in ALS (Luisier *et al.,* 2018). To this end we analysed data from a method called individual-nucleotide resolution cross-linking and immunoprecipitation (iCLIP), which allows identification of the RNA binding targets of the FUS protein. Using this approach, we found that the FUS protein binds extensively to the aberrantly retained intron 9 within the SFPQ transcript (Figure 3A&B), which we identified as the most significantly retained intron across diverse ALS mutations (Luisier *et al.,* 2018). We next confirmed that the SFPQ intron-retaining transcript is exported to the cytoplasm using nuclear-cytoplasmic cellular fractionation and qPCR (Figure 3C), showing an increased proportion of SFPQ intron retaining transcript in the cytosol of VCP mutant cultures. We employed single molecule RNA fluorescence in situ hybridization (smFISH) as orthogonal validation that this transcript is exported from the nucleus (Figure 3D).

**Figure 3.**
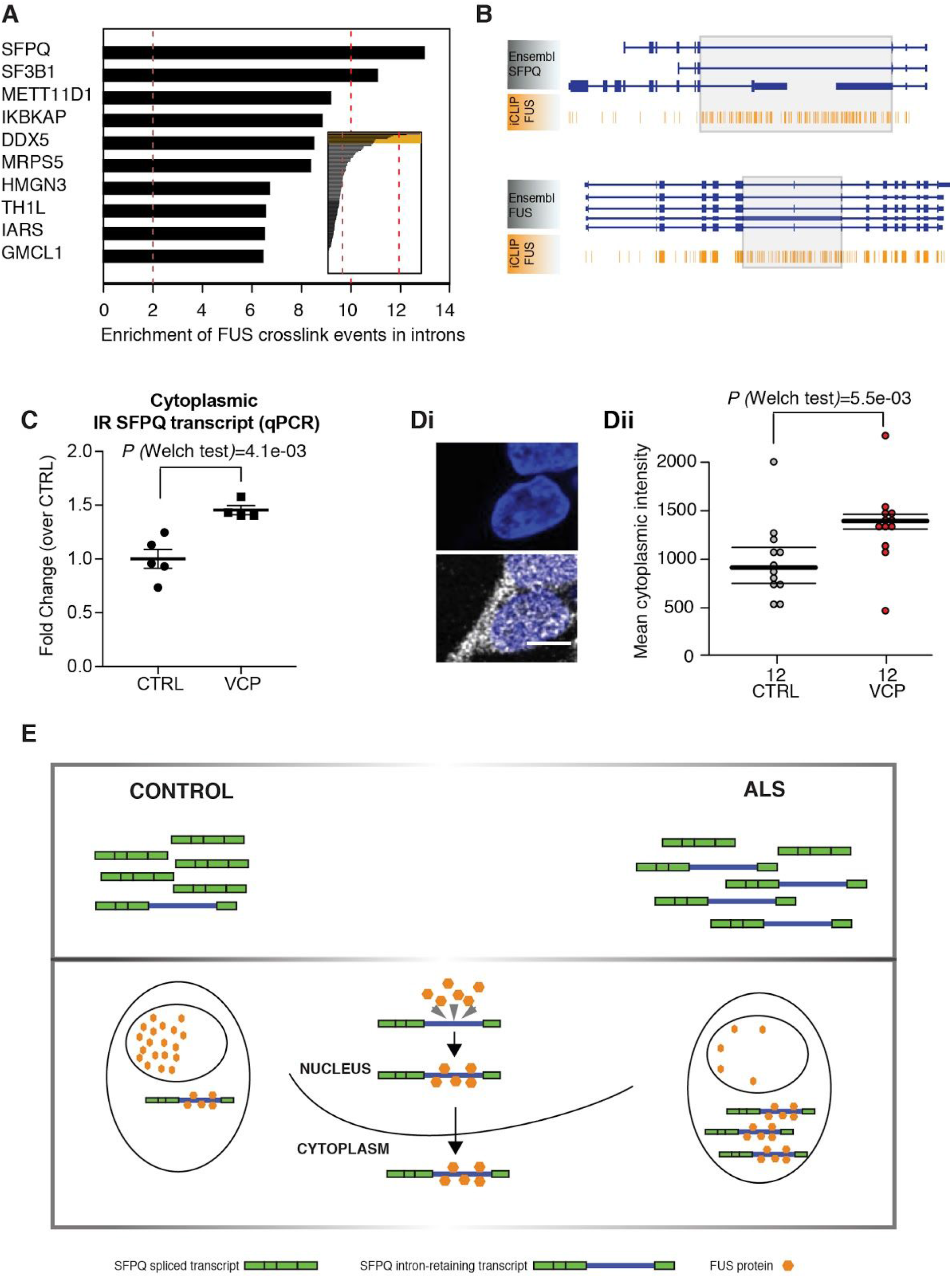
FUS binds to an aberrantly retained intron within the SFPQ transcript, which is exported from the nucleus. **(A)** Bar plots depicting the level of enrichment in FUS-binding to the retained intron compared with the non-retained introns within the same gene. Analysis of the 167 previously reported as aberrantly retained introns in ALS. **(B)** Genome browser view of FUS iCLIP crosslinking events along the SFPQ and FUS transcript. Grey boxes highlight the location of retained introns. **(C)** Dot plot showing the levels of cytoplasmic IR SFPQ transcripts measured by qPCR. IR transcripts level were measured using an IR specific primer pair and normalised over the expression levels of a constitutive SFPQ exon, as previously described (Luisier *et al.,* 2018). N= 5 control lines and N=4 VCP lines, mean ± s.e.m. from two independent replicates per line, each analysed in technical duplicate. P=0.0041 from Welch’s test. **(D)** i) Single molecule fluorescence in situ hybridisation (smFISH) showing SFPQ intron-retaining transcript in the cytoplasm. The upper panel shows the nuclear staining only for context; the probe clearly detects intron-retaining transcripts in the cytoplasm. ii) Quantification and statistical analysis of smFISH performed on 3 control lines and 3 VCP mutant lines using Welch’s test. **(E)** Schematic diagram of proposed model. Cartoon summarising the functional consequence of aberrant IR in SFPQ transcript. The 9kb intron 8/9 in SFPQ is retained and exported to the cytoplasm. The FUS protein binds extensively to this retained intron. The FUS protein itself is abundantly relocalised from the nucleus to the cytoplasm in diverse ALS models (human iPSC and mouse transgenic), as well as post-mortem samples from sporadic ALS patients.

## Discussion

The core finding of our study is that the FUS protein is mislocalised from the nucleus to the cytoplasm in cases beyond FUS mutation-related ALS. Specifically, we find FUS mislocalisation in VCP mutation-related ALS and, crucially, in sporadic ALS. Our findings have been carefully cross-validated in iPSC lines (4 VCP mutant and 4 control iPSC lines), a mouse transgenic model (3 VCP mutant and 3 control mice), and post-mortem tissue (12 sporadic ALS cases and 8 control cases). The pervasive mislocalisation of FUS has likely evaded detection thus far as FUS largely remains unaggregated in the cytoplasm, rather than forming part of the TDP-43 aggregates in sporadic ALS cases.

Our findings support a model whereby FUS mislocalisation from the nucleus to the cytoplasm occurs in the majority of ALS cases, but it generally does not appear to aggregate in the cytoplasm. Nuclear loss of FUS protein may impair pre-mRNA splicing whilst the possibility of a cytosolic toxic gain of function is also noteworthy in light of recent studies (López-Erauskin *et al.,* 2018). Interestingly, TDP-43 aggregation is also observed in most familial and sporadic ALS cases, except from those caused by SOD1 mutations. Thus, FUS mislocalisation appears to occur in a similar majority of cases to TDP-43 aggregation. Importantly, the mislocalised FUS is observed in mouse and iPSC models at early stages, before the onset of TDP-43 aggregates, indicating that the mislocalised unaggregated FUS might play a causative role early in the ALS disease process, perhaps by creating a more aggregation-prone cytoplasmic environment.

Our analysis of iCLIP data suggest that FUS binds avidly to the aberrantly retained intron of the SFPQ transcript in ALS. Cumulatively, our data are consistent with a working hypothesis that wild-type FUS might travel out of the nucleus when bound to the aberrantly retained intron 9 of the SFPQ transcript in ALS (Figure 3E), though confirmation of such a mechanism will require detailed molecular follow-up work. The lack of evidence for FUS mislocalisation in SOD1 mutant MNs despite the presence of SFPQ intron retention (Luisier *et al.,* 2018) suggests a mutation-dependent variation in the repertoire of RBPs that engage intron-retaining transcripts. SFPQ protein can be sequestered by the SFPQ intron-retaining transcript across the different mutations, whereas in the context of SOD1 mutations, FUS might not be significantly sequestered and/or exported in this manner.

In summary we report a previously unrecognized widespread mislocalisation (but not aggregation) of FUS in ALS, and propose a putative context-specific mechanism for this through its interaction with the ALS-related aberrantly retained intron 9 in SFPQ. These findings raise the prospect of targeting the nuclear mislocalistion of unaggregated FUS as a putative therapeutic strategy in ALS.

## Methods

### iPSC culture and motor neuron differentiation

iPSC were maintained using standard protocols. Motor neuron differentiation was carried out as *previously* described (Hall *et al.,* 2017; Luisier *et al.,* 2018). See Supplementary Methods for more detailed information. Details of iPSC lines are provided in Table S1.

### Animals, transgenic models and tissue processing

All animal experiments described in this study were carried out under license from the UK Home Office, and were approved by the Ethical Review Panel of the Institute of Neurology. The following transgenic mouse lines were used: (1) *SOD1^G93A^*mice (*B6SJL-Tg(SOD1*G93A)1Gur/J,* Jackson Laboratories) (2) over-expressing mutant human VCPA232E were generated by J. Paul Taylor et al, St Jude Children’s Research Hospital, Memphis, TN, USA and are described in (Custer *et al.,* 2010) and bred to wild-type C57 Black 6 (*C57/B6*) background. Wild type *C56BL/6-SJL* mixed background (Jackson Laboratories) were used as control. For tissue collection, animals were injected with terminal anaesthesia (pentobarbital sodium, Euthatal) and were transcardially perfused with 4% paraformaldehyde. The lumbar region of the spinal cord was removed and post-fixed with 4% paraformaldehyde and cryoprotected overnight with 30% sucrose; 10 or 20μm serial transverse cryosections were cut for immunofluorescence staining.

### Human post mortem tissue

Snap frozen tissue sections derived from lumbar spinal cords of three age and sex matched sporadic ALS patients (see Table S2). Death to snap-freezing delay times were also comparable between the groups (delay: 30.13±12.87 and 27.75±10.63 hours, for control and sALS patients respectively). The spinal cord samples were obtained from the tissue bank NeuroResource, UCL Institute of Neurology, London, UK. Samples were donated to the tissue bank with written tissue donor informed consent following ethical review by the NHS NRES Committee London - Central and stored under a Research Sector Licence from the UK Human Tissue Authority (HTA).

### Immunolabelling and imaging

For immunocytochemistry and immunohistochemistry, samples were blocked in 10% normal goat serum (NGS) or 10% normal donkey serum (NDS) as appropriate and permeabilised in 0.3% Triton X-100 (Sigma-Aldrich; in PBS) at room temperature (RT) for 1 hour. Immunolabelling was performed with primary antibodies in NGS (5%) and Triton X-100 (0.1% in PBS) at 4°C overnight followed by species-specific secondary antibodies for 1 hour at RT and DAPI nuclear counterstain (100 ng/ml) for 10 min at RT. For human post-mortem samples fixation and permeabilisation in cold methanol (−20°C, 20 minutes) was performed before the immunostaining. Primary antibodies were diluted as follows: goat anti ChAT (Millipore, AB144P) 1:100; rabbit and mouse anti FUS (Abcam, ab84078 1:500; and Santa Cruz, sc-47711 1:100 respectively). Images were acquired using either a 710 Laser Scanning Confocal Microscope (Zeiss) or the Opera Phenix High-Content Screening System (Perkin Elmer). Images were acquired as confocal z-stacks with a z step of 1μm and processed to obtain a maximum intensity projection. For the analysis of nuclear/cytoplasmic ratio of FUS in iPSC-derived cells, images were analysed using the Columbus Image Analysis System (Perkin Elmer). The post-mortem tissue sections were analysed using Fiji. MNs were identified by ChAT immunoreactivity, and the nuclear and cytoplasmic area were manually drawn based on DAPI and ChAT staining respectively. For each region of interest, the average FUS immunoreactivity intensity was measured, background was subtracted, and the ratio between nuclear and cytoplasmic average intensity was calculated.

### Cell fractionation, RNA extraction, reverse transcription and qPCR

For the biochemical fractionation of iPSC-derived NPCs the Ambion PARIS kit (Thermo Fisher Scientific) was used following manufacturer’s instructions. A cytosolic fraction was obtained by lysing the cultures for 10 minutes in ice-cold Cell Fractionation buffer. Nuclei were lysed in 8M Urea Nuclear Lysis Buffer, containing 50mM Tris-HCl (pH 8), 100mM NaCl, 0.1% SDS, 1mM DTT. Both lysis buffer contained 0.1 U/μl RiboLock RNase Inhibitor (Thermo Fisher Scientific). RNA was extracted from both fractions using the Promega Maxwell RSC simplyRNA cells kit including DNase treatment, alongside the Maxwell RSC instrument. Reverse transcription was performed using the Revert Aid First Strand cDNA Synthesis Kit (ThermoFisher Scientific) using 1μg of RNA and random hexamers. qPCR was performed using the PowerUP SYBR Green Master Mix (ThermoFisher Scientific) and the the QuantStudio 6 Flex Real-Time PCR System (Applied Biosystems). Specific amplification was determined by melt curve analysis and agarose gel electrophoresis of the PCR products. Analysis of intron retaining transcripts was performed as previously described (Luisier *et al.,* 2018). Levels of IR (primer pair F2R2) were normalised over the expression level of each individual gene (primer pair F1R1). Primers user were as follows. SFPQ F1: GCCGAATGGGCTACATGGAT. SFPQ R1: TCAGTACGCATGTCACTTCCC. SFPQ F2: GTGGATCGACTCATTGGTGA. SFPQ R2: TTCCTCTAGGACCCTGTCCA.

### Single molecule FISH

Based on (Raj *et al.,* 2008), iPSC-derived NPCs (day 7 after beginning of neural conversion) that were grown on Geltrex coated sterile 2-well μ-Slides (80286, Ibidi) were washed twice for 5 min with 1x PBS, and fixed using 4 % methanol-free formaldehyde (10321714, Thermo Fisher Scientific) for 10 min at RT. Following additional two wash steps, cells were permeabilized using 70 % ethanol for 12 h at 4 °C, and were washed twice again. Then preparations were incubated for 15 min with hybridization buffer prepared using 2x saline-sodium citrate (SSC) solution / 10 % deionized formamide (4610-OP, Calbiochem). Hybridization with Stellaris FISH probes was done in a total volume of 50 μl hybridization buffer containing 50 μg competitor tRNA from E.coli (10109541001, Roche Diagnostics), 10 % Dextran Sulfate (9011-18-1, VWR), 2 mg/ml UltraPure BSA (AM2616, Thermo Fisher Scientific), and 10 mM vanadyl-ribonucleoside complex (S1402S, New England BioLabs) with probes at final concentration of 1 ng/μl. Preparations were covered with parafilm and incubated at 37 °C for 5 h, and afterwards washed twice with pre-warmed 2xSCC/10 % formamide for 30 min at 37 °C. Finally the preparations were washed twice with 1x PBS at RT, and then mounted using 10 μl ProLong Gold Antifade Reagent containing DAPI (9071S, New England Biolabs). The slides were imaged when the mounting medium was fully cured >12 h.

Probes were designed using the Probe Designer software from Biosearch Technologies and were provided by same vendor. Probes included were designed against SFPQ intron conjugated to Quasar^®^570 (SMF-2037-1) and mature SFPQ conjugated to Quasar^®^670 (sequences of probes available upon request). Quantification of hybridization signal was performed using custom spot-intensity detection algorithm in DAPI segmented cells to separate nuclear and cytoplasmic signal.

### Data analysis of FUS protein sub-cellular localization

We used R (R Core Team, 2012) and lme4 (Bates, Maechler & Bolker, 2012) to perform linear mixed effects analysis of the relationship between FUS localisation and VCP or SOD1 mutation, as well as with sporadic ALS, that accounts for idiosyncratic variation due to either animal or individual differences. As fixed effects, we either entered the mutation or the ALS disease variable into the model. As random effects, we had intercepts for either animals (SOD1, VCP) or patients and batches. Visual inspection of residual plots did not reveal any obvious deviations from homoscedasticity or normality. P-values were obtained by likelihood ratio tests of the full model with the effect in question against the model without the effect in question.

### Mapping of iCLIP data

Raw FUS iCLIP data can be accessed via https://imaps.genialis.com/ (Attig *et al.,* 2018). Before alignment, two-stage adapter removal was performed using Cutadapt according to the ENCODE iCLIP standard operating procedure. A two-stage approach was also used for alignment. First, Bowtie2 was used to remove reads aligning to rRNA or tRNA. Then, STAR was used to align the remaining reads to GRCh38, with only uniquely mapping reads retained. PCR duplicates were collapsed based on the unique molecular identifiers and mapping locations. The nucleotide-resolution crosslink position was calculated as the coordinate immediately preceding the reverse transcription truncation event.

## Compliance with ethical standards

For human iPSC work: Informed consent was obtained from all patients and healthy controls in this study. Experimental protocols were all carried out according to approved regulations and guidelines by UCLH’s National Hospital for Neurology and Neurosurgery and UCL Queen Square Institute of Neurology joint research ethics committee (09/0272). The human post-mortem spinal cord samples were obtained from the tissue bank NeuroResource, UCL Queen Square Institute of Neurology, London, UK. Samples were donated to the tissue bank with written tissue donor informed consent following ethical review by the NHS NRES Committee London - Central and stored under a Research Sector Licence from the UK Human Tissue Authority (HTA). All animal experiments described in this study were carried out under licence from the UK Home Office, and were approved by the Ethical Review Panel of the Institute of Neurology.

## Acknowledgments

The authors wish to thank Martina Hallegger and Roberto Simone for their input regarding primer design, Anob M Chakrabarti for sharing BED files of aligned iCLIP data. We thank Benjamin Clarke, Mhoriam Ahmed and members of the Schiavo and Greensmith Labs for valuable technical support. This work was supported by the Francis Crick Institute which receives its core funding from Cancer Research UK (FC010110), the UK Medical Research Council (FC010110), and the Wellcome Trust (FC010110). R.P. holds an MRC Senior Clinical Fellowship [MR/S006591/1]. N.M.L. and J.U. are supported by a Wellcome Trust Senior Investigator Award [103760/Z/14/Z], an MRC eMedLab Medical Bioinformatics Infrastructure Award to N.M.L. (MR/L016311/1), the UCL Grand Challenges Award (C.E.H.), a Marie Curie Post-doctoral Research Fellowship (657749-NeuroUTR) and an Advanced Postdoc Mobility Fellowship from the Swiss National Science Foundation (P300PA_174461) to R.L. N.M.L. is a Winton Group Leader in recognition of the Winton Charitable Foundation’s support towards the establishment of the Francis Crick Institute.

## Supplementary information

### Supplementary methods

#### Derivation of Human Fibroblasts and iPSC

Dermal fibroblasts were cultured in OptiMEM +10% FCS medium. The following episomal plasmids were transfected for iPSC generation: pCXLE hOct4 shp53, pCXLE hSK, and pCXLE hUL (Addgene), as previously reported (Okita *et al.,* 2011). Details of the lines used in this study are provided in Table S1. Three of the control lines used (control 2 and control 3, and control 5) are commercially available and were purchased from Coriell (cat. number ND41866*C), ThermoFisher Scientific (cat. number A18945) and Cedars Sinai (Cat.number CS02iCTR-NTn4) respectively.

#### Cell Culture and motor neuron differentiation

Induced PSCs were maintained on Geltrex (Life Technologies) with Essential 8 Medium media (Life Technologies), and passaged using EDTA (Life Technologies, 0.5mM). All cell cultures were maintained at 37°C and 5% carbon dioxide. For motor neuron (MN) differentiation, iPSCs were first differentiated to neuroepithelium by plating to 100% confluency in chemically defined medium consisting of DMEM/F12 Glutamax, Neurobasal, L-Glutamine, N2 supplement, non-essential amino acids, B27 supplement, β-mercaptoethanol (all from Life Technologies) and insulin (Sigma). Treatment with small molecules from day 0-7 was as follows: 1μM Dorsomorphin (Millipore), 2μM SB431542 (Tocris Bioscience), and 3.3μM CHIR99021 (Miltenyi Biotec). At day 8, cells patterned for 7 days with 0.5μM retinoic acid and 1μM Purmorphamine. At day 14 spinal cord MN precursors were treated with 0.1μM Purmorphamine for a further 4 days before being terminally differentiated for >10 days in 0.1 μM Compound E (Enzo Life Sciences) to promote cell cycle exit. Throughout the neural conversion and patterning phase (D0-18) the neuroepithelial layer was enzymatically dissociated twice (at D4-5 and D10-12) using dispase (GIBCO, 1 mg ml^-1^). At relevant timepoints cells were harvested for RNA extraction or fixed in 4% paraformaldehyde for immunolabelling.

**Supplementary table 1.** Details of iPSC lines used in this study.

| iPSC line | Mutation present | Age of Donor | Age at disease onset | Sex of Donor |
| --- | --- | --- | --- | --- |
| **CTRL 1** | None | 78 | N/A | Male |
| **CTRL 2** | None | 64 | N/A | Male |
| **CTRL 3** | None | Unknown | N/A | Female |
| **CTRL 4** | None | 51 | N/A | Female |
| **CTRL 5** | None | 51 | N/A | Male |
| **MUT 1** | VCP R155C | 43 | 40 | Female |
| **MUT 2** | VCP R155C | 43 | 40 | Female |
| **MUT 3** | VCP R191Q | 42 | 36 | Male |
| **MUT 4** | VCP R191Q | 42 | 36 | Male |

**Table S2.** Information of the patients used in the post mortem tissue analysis.

| **Age** | **Gender** | **Category** | **Cause of death** | **Post mortem delay (hours)** |
| --- | --- | --- | --- | --- |
| 71 | M | Control | Burst aortic aneurysm. | 25 |
| 68 | F | Control | Colorectal metastatic cancer | 23 |
| 68 | M | Control | Heart disease | 40 |
| 70 | M | Control | Bronchopneumonia and heart failure | 53 |
| 77 | M | Control | Pulmonary fibrosis | 32 |
| 80 | F | Control | Pulmonary embolism | 24 |
| 81 | M | Control | Lung carcinoma; bilateral bronchopneumonia | 10 |
| 67 | F | Control | Bronchopneumonia | 34 |
| 69 | M | sALS | MND & stage 2 respiratory failure. | 19 |
| 61 | M | sALS | MND. | 29 |
| 74 | F | sALS | MND. | 27 |
| 70 | M | sALS | MND. | 40 |
| 81 | F | sALS | MND. | 33 |
| 65 | F | sALS | MND | 30 |
| 70 | F | sALS | MND | 26 |
| 58 | M | sALS | MND | 49 |
| 69 | M | sALS | MND | 19 |
| 66 | F | sALS | MND | 15 |
| 77 | F | sALS | Bronchopneumonia and MND | 34 |
| 80 | F | sALS | MND | 12 |

